# Disentanglement of Resting State Brain Networks for Localizing Epileptogenic Zone in Focal Epilepsy

**DOI:** 10.1101/2022.06.13.495945

**Authors:** Shuai Ye, Anto Bagić, Bin He

## Abstract

Resting state electromagnetic recordings have been analyzed in epilepsy patients aiding presurgical evaluation. However, it has been rarely explored how pathological networks can be separated and thus used for epileptogenic focus localization purpose. We proposed here a resting state EEG/MEG analysis framework, to disentangle brain functional networks represented by electrophysiological oscillations. Firstly, by using an Embedded Hidden Markov Model (EHMM), we constructed a state space for resting state recordings consisting of brain states with different spatiotemporal patterns. After that, functional connectivity analysis along with graph theory were applied on the extracted brain states to quantify the network features of the extracted brain states, and we determine the source location of pathological states based on these features. The EHMM model was rigorously evaluated using computer simulations. Our simulation results revealed the proposed framework can extract brain states with high accuracy regarding both spatial and temporal profiles. We than validated the entire framework as compared with clinical ground truth in 10 patients with drug-resistant focal epilepsy who underwent MEG recordings. We segmented the resting state MEG recordings into a few brain states with diverse connectivity patterns and extracted pathological brain states by applying graph theory on the constructed functional networks. We showed reasonable localization results using the extracted pathological brain states in 6/10 patients, as compared to the invasive clinical findings. The framework can serve as an objective tool in extracting brain functional networks from noninvasive resting state electromagnetic recordings. It promises to aid presurgical evaluation guiding intracranial EEG electrodes implantation.

## Introduction

Brain function and dysfunction are encoded in spatiotemporally distributed networks. Functional neuroimaging has evolved from activation imaging, such as functional MRI task-based activation imaging and EEG/MEG (E/MEG) source localization of focal activities, to distributed imaging of brain networks, such as resting state fMRI connectivity imaging (Boerwinkle et al., 2017; Khoo et al., 2017; Lee et al., 2018), and resting state spontaneous mapping and imaging from electrophysiological measurements (Canuet et al., 2011; Case et al., 2018; Coito et al., 2016; Mantini et al., 2007). Recent studies suggest that network analysis of resting state intracranial EEG recordings can allow localization of seizure-onset-zone (SOZ) for up to 88% accuracy and predict seizure outcome (Jiang et al., 2022). A scientific question of importance to neuroimaging is: can we estimate brain activity and connectivity associated with tasks and behaviors from resting state electrophysiological recordings over the scalp? The answer to this question may have implications to broad neuroimaging of brain function and dysfunction. In particular, this may have a significant impact to identification and imaging of epileptic networks from brief resting state scalp E/MEG recordings.

Epilepsy is a chronic neurological disorder affecting over 65 million people in the world and 3.5 million people in the United State alone (Moshé et al., 2015; Thijs et al., 2019). While antiseizure medication (ASMs) serve as the primary treatment, about 30% of the patients do not respond to any combination of multiple ASMs (Engel, 2008; Palmini et al., 1991). These patients may benefit from surgical intervention (Duncan et al., 2016; Rosenow and Lüders, 2001) or brain stimulation implants (Ben-Menachem, 2002; Vonck et al., 2002) targeting at the epileptogenic zone (EZ) (Rosenow and Lüders, 2001). Thus, accurate estimation of the epileptic network and localization of the EZ plays a critically important role in guiding the resection surgery and in directing electrode implants for brain stimulation.

Clinically, the criteria for defining the EZ to cure the patients are far from being standardized (Jehi, 2018; Stefan et al., 1987). In practice, patients go through a comprehensive presurgical evaluation including multiple imaging modalities. Among all functional imaging approaches, the high temporal resolution of E/MEG allows the detection of the spatiotemporal dynamics more efficiently than other non-invasive imaging techniques (Agirre-Arrizubieta et al., 2009; Cohen, 1968; Hara et al., 2007; Pizzo et al., 2019; Sekihara et al., 2001; Sohrabpour and He, 2021). To further improve the spatial resolution of E/MEG and enable exploration of cortical activities, electrophysiological source imaging (ESI) (Brodbeck et al., 2011; He et al., 2020, 2018; Michel and He, 2017) is adopted to localize and image the sources of scalp recorded E/MEG signals. The localization accuracy of ESI in presurgical evaluation has been demonstrated by numerous studies and has emerged as a viable option for guiding surgical interventions (Brodbeck et al., 2011; Kaiboriboon et al., 2012).

To better estimate the epileptic network, many studies with E/MEG recordings have utilized curated abnormal signals such as seizures or interictal epileptic discharges (IEDs) (Agirre-Arrizubieta et al., 2009; Chowdhury et al., 2015; de Curtis et al., 2013; Ding and He, 2006; Gotman and Gloor, 1976; Gotman and Marciani, 1985; He et al., 1987; Heers et al., 2016, 2014; Sohrabpour et al., 2016, 2020; Yang et al., 2011; Ye et al., 2021) to localize the EZ. While the convenience of analyzing selected IEDs must be acknowledged, it is also widely known that the selection of IEDs is highly dependent on the readers’ expertise (Kural et al., 2020; Ye et al., 2021), not to mention the extensive human labor involved during this procedure. Co-existing of multiple types of IED also adds to the burden of visual inspection and identification. Seizures, while showing superior accuracy of localizing Seizure Onset Zone (SOZ) with advanced analysis techniques (Lu et al., 2012; Yang et al., 2011; Ye et al., 2021), remains challenging to capture in many clinical routine examinations. More approachable and data-driven methodologies that can be integrated into clinical decision-making are still in great need.

Interests in utilizing the resting state have been emerging, whose major advantage is that they can be estimated without the need to wait for a seizure to occur or a concordant IED to be identified (Hsiao et al., 2015; Krishnan et al., 2015). Recent discoveries using fMRI suggested a limited number of large-scale distributed networks of temporally correlated spontaneous activity (Beckmann et al., 2005; Boerwinkle et al., 2017; Damoiseaux et al., 2006; Smith et al., 2013; Zhang et al., 2015), while the task-positive cerebral activities may be highly complicated. In E/MEG signal with high temporal resolution, similar networks have been extracted as transient activations and shown to have distinct band-limited oscillatory power (Baker et al., 2014; Vidaurre et al., 2016). Altered dynamics of these large-scale networks, on the other hand, has been associated with various neurological disorders such as Alzheimer’s disease (De Haan et al., 2008; Stam et al., 2005), Parkinson’s disease (Cao et al., 2015; Müller et al., 2001), chronic pain (Case et al., 2019)and schizophrenia (Kottaram et al., 2019; Lefebvre et al., 2016; Naim-Feil et al., 2018). These types of large-scale network analysis are of greater interest for epilepsy patients due to the relationship between EZ localization and epileptic network estimation, where it is essential to identify and then remove or disconnect certain nodes of the epileptic network. In other words, disentanglement of pathological network for the purpose of source localization is of importance for focal epilepsy patients, and potentially for delineating brain normal function as well as various dysfunctions.

In this context, one natural direction is to consider the E/MEG resting state signal as a finite-state model that can be described in terms of intrinsic spatiotemporal dynamics (Khanna et al., 2015). The method of studying E/MEG resting state recordings, therefore, is by defining the state space based on variables of interest and describing changes in brain activity in terms of state characteristics, such as the duration or frequency of occurrence of specific states. One realization of the method is the so-called “microstates”, where four discrete states are defined by E/MEG topography and remains stable for 80-120 ms before rapidly transitioning to another ((Lehmann et al., 1987; Michel and Koenig, 2018; Pascual-Marqui et al., 1995) Several studies following this framework have explored the differences between patients with neurological disorders and healthy controls, where significant differences were found in the occurrence of states or the transition matrix (Khanna et al., 2015; Piorecka et al., 2018). While such analysis confirmed the existence of underlying abnormal pattern with or without IED existing(Ahmadi et al., 2020; Khanna et al., 2015; Yuan et al., 2012), abundant spatiotemporal information is missed during the procedure since the signal is filtered into certain frequency bands (alpha, theta, etc.) before being fed into the framework (Lehmann et al., 1987; Michel and Koenig, 2018; Poulsen et al., 2018). Moreover, the brain state extraction is based on four topography templates, which makes it challenging to obtain foci-related localization information (Shaw et al., 2019).

Hidden Markov Model (HMM) was adopted to address the issues and has shown comparable outcome if used in extracting the four aforementioned microstates (Eddy, 2004; Lee and Choi, 2003; Ossadtchi et al., 2005). In HMM, each state will be characterized with the transition to other states, instead of relying on the topographic templates, thus more refined classification of brain states is achievable (Rabiner and Juang, 1986). To further address the spatiotemporal information as missed by static state-space analysis, one way is to explicitly model the spatiotemporal correlation between channels in a time window, which was used in previous studies when a limited number of channels is involved (Quinn et al., 2018; Sohrabpour et al., 2016). Alternatively, a computationally feasible solution is through space embedding (Vidaurre et al., 2016). Embedded Hidden Markov Model (EHMM) has been shown to effectively extract hidden brain states in source-space as applied in event related potential analysis or spontaneous activities in healthy subjects (Jebara et al., 2007; Quinn et al., 2018; Vidaurre et al., 2018). These extracted brain states were compared to functional brain networks from functional magnetic resonance imaging (fMRI) or PET studies, showing that spontaneous brain activities are not random but forming highly coherent functional networks, which can be separated with the EHMM framework (Vidaurre et al., 2016).

The existing evidence suggests that the transient neural activity of the brain has quasistable properties and could be evaluated with a network view. To further quantify brain dynamics, network analyses from various structural and functional modalities have also emerged aiming at providing a better understanding of network characteristics of the epileptic brain, especially the changes in these characteristics related to surgery outcome. A network measurement, betweenness centrality, was found to correlate with the location of the resected cortical regions in patients who were seizure free after surgery (Wilke et al., 2011). Small-worldness was also found to be an important property where brain networks could be distinguished from random networks and was used to differentiate epilepsy patients from controls (Farahani et al., 2019; Lithari et al., 2012; Smit et al., 2008). Several other studies (Coito et al., 2016; Elisevich et al., 2011) have showed disruptions in various network features, and achieved lateralization in temporal lobe epilepsy patients. In this regard, network measures provide a quantitative approach to characterize complex network dynamics and can lead to a better understanding of the epileptic network in addition to providing a valuable diagnostic tool.

In this work, we aim to integrate the strength of brain state space approaches and network analysis based on connectivity, to better disentangle the pathological networks from resting state E/MEG signal, thus, to estimate the EZ. We firstly adopted EHMM as the brain state extraction algorithm and verified its accuracy with computer simulations. We then applied the framework to MEG recordings of focal epilepsy patients and analyzed the extracted states with commonly used connectivity metrics to identify the pathological states. By doing so, we observed concordant results between the estimated foci from the proposed framework and the seizure onset zone identified from clinical evaluations, especially in those patients who had abundant IEDs.

## Methods

The general analysis framework can be summarized as a two-step strategy as shown in Figure 1. Firstly, we designed a brain state extraction framework by adapting an Embedded Hidden Markov Model and verified the model using Monte Carlo simulation. Next, the extracted brain states underwent a connectivity analysis to extract the connectivity features and related network attributes. A Graph Feature Index was designed based on the network features, to identify the pathological states among all extracted brain states. The extracted pathological states were compared to the clinical findings to evaluate the accuracy of the proposed framework.

**Figure 1.**
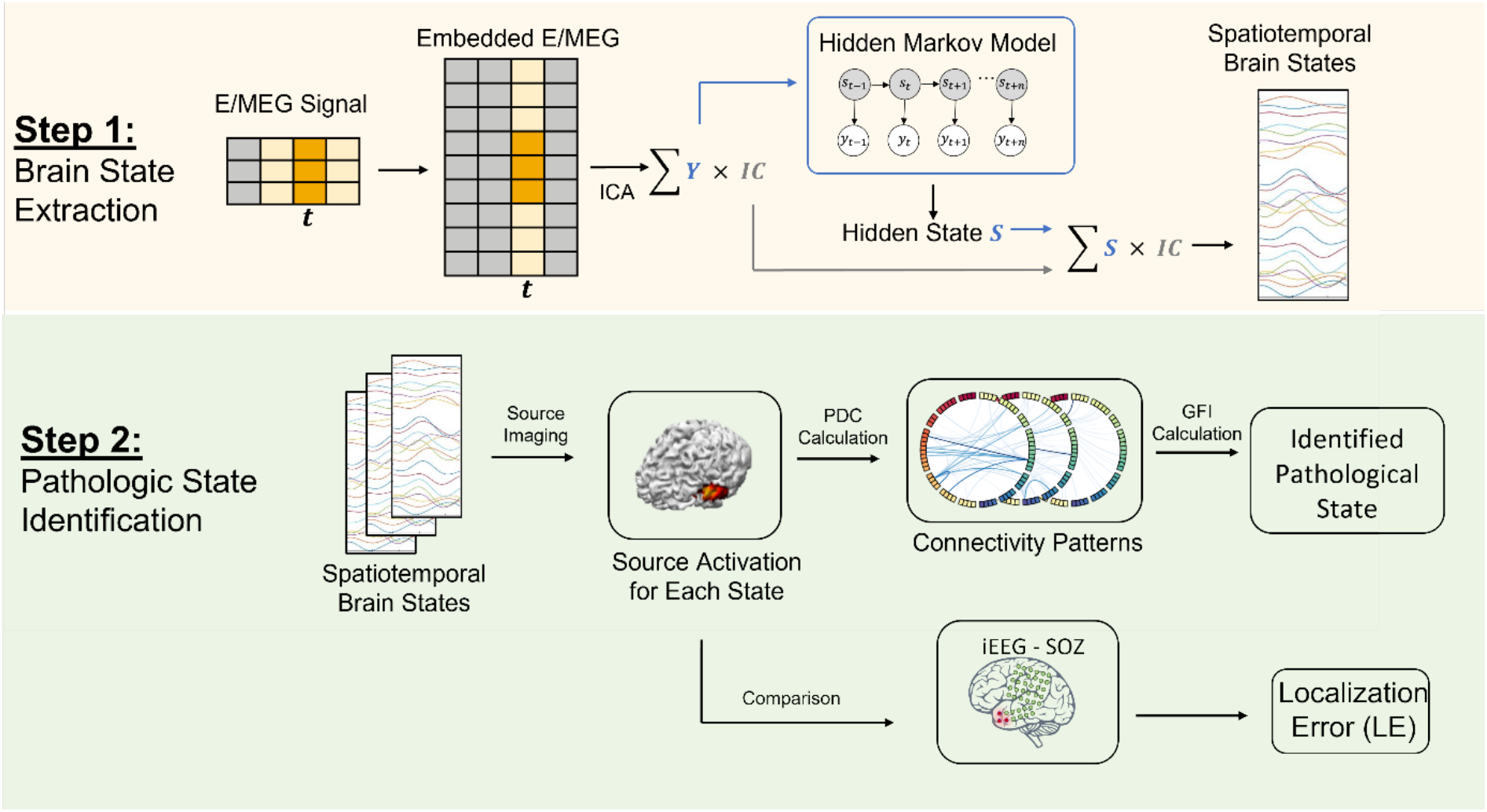
The overall study pipeline. i) In step 1, a time window of the original E/MEG data were embedded to create the embedded E/MEG. Independent Component Analysis (ICA) was applied on the data to decompose the data into topographical components and their corresponding activation. The activations were then put through the Hidden Markov Model to obtain the brain states. The estimated states were multiplied by the IC again to transform back to the original E/MEG space to generate the spatiotemporal brain states. ii). In step 2, source imaging was applied on the extracted brain states to project sensor space activity to the source space activity. The connectivity features were extracted using Partial Directed Coherence (PDC) and were further used to identify the pathological states. The localization error of the identified pathological states was used to evaluate the accuracy of the proposed framework.

### Embedded Hidden Markov Model

In the HMM, it is assumed that a time series can be described using a hidden sequence of a finite number of states and each state will be characterized with the transition to other states. To facilitate the process, we adapted a well-established Bayesian variational learning algorithm. Assuming state space dimension has the dimension of *K*, hidden state variables s = {*s*_1_…*s_T_*}, and the observed data is denoted by y = {*y*_1_ …*y_T_*} for the time points *t*_1_ …*t_T_*. The initial state probability is denoted by π = {*π*_1_ …*π_K_*}, where *π*_i_ = *p*(*s*_1_ = *q*_i_) where variable *q*_i_ represent the ith state for i=1,…, K. Set *a_t_* to be the transition probability where element *α_t,(i,j)_* = *p*(*s*_*t*+1_ = *q_j_*|*s_t_* = *q_i_*) describes the probability of transitioning from hidden state *s*_i_ to *s*_j_ from time *t* to (*t* + 1). Let *A* = {*A*_1_, …,*A*_T_}, where *A*_t_ denote the transition probability matrix at time *t*.

We then set the term *b_i_*(*y_t_*) = *p*(*y_t_*|*s_t_* = *q*_i_, *θ*) to demote the emission probability distribution. In order to describe the multi-channel E/MEG data, we assume that the emission probability for hidden state as the multivariate Gaussian distribution described by

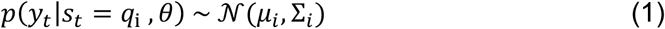

where *θ_i_* = {*μ_i_*, Σ_*i*_, *μ_i_* being the mean vector, and Σ_*i*_ being the covariance matrix. We then denote the parameters as Θ = {*θ*_1_, …,*θ_K_*}.

With all the parameters, the joint distribution p(y, s|Θ,*A,π*) is then written as

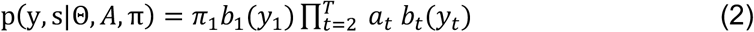

In the basic HMM, the observation y should be the original E/MEG data. This model, on one hand, detects the instantaneous spatial changes and can trace state transitions, but on the other hand, discards rich information of the spectral interactions among the channels (e.g., oscillations leading to repeating patterns, propagation of the oscillations, etc.). EHMM can be applied to capture temporal dynamics of the signal. In this study, we adapted the framework of a time-delay embedded HMM (Vidaurre et al., 2018), where the E/MEG activities over a time window are described using a Gaussian distribution with zero means which is equivalent to using a standard HMM on an embedding transformation of the original data. Many methods can be applied to construct the embedding space of the original data (Jebara et al., 2007; Seide et al., 2003; Vidaurre et al., 2016). Naturally, Independent Component Analysis (ICA) is one of the top choices, as the goal of ICA (making the outputs statistically independent being sensitive to higher-order statistics) aligns with the features of data to be extracted, as shown by numerous previous studies (Chen et al., 2013; Hsu et al., 2018; Nam et al., 2002; Patel et al., 2008; Seide et al., 2003; Ye et al., 2021; Yuan et al., 2012).

With ICA as an embedding tool, instead of feeding the entire embedded E/MEG into the HMM model, we firstly performed an ICA decomposition on the embedded E/MEG and the original signal can be written as:

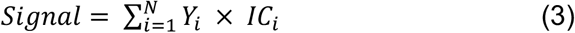

where *IC_i_* is the independent components, and *Y_i_* being the activation for the components. Thus, we transformed the original signal from actual channel space (each value representing the activation amplitude in one E/MEG channel) into the component space (each value representing the activation amplitude for one independent component). Instead of the actual signal, *Y* was used as the input to the Hidden Markov Model. After HMM inference, the hidden states are extracted and represented as *S_j_*, which is a weighting with dimension of *N* defined on the component space.

We then multiply *S_j_* to the components as:

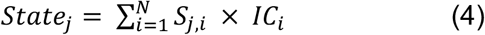

to reverse from component space to the actual data space. Eventually, we have the extracted states as transient E/MEG segments representing brain states inferred from the Hidden Markov Model.

Variational Bayes (VB) were used to infer the model parameters, which assumes additional factorizations in the space of parameters and needs all prior distributions to be conjugate (Quinn et al., 2018; Vidaurre et al., 2018, 2016). By using the Expectation– Maximization algorithm acting on one group of parameters at a time, variational Bayes inference minimizes the so-called free energy (Rezek and Roberts, 2005).

The code regarding HMM was written in MATLAB 2019b and partial code regarding the free energy evaluation is based on the literatures and toolboxes provided (Quinn et al., 2018; Rezek and Roberts, 2005; Vidaurre et al., 2018, 2016).

### Simulation Protocol

Monte Carlo simulation with synthesized MEG data was used to verify if the EHMM model can extract states accurately. Synthesized MEG data was generated based on the following assumptions: i) the brain states can be segmented into a few distinguished states; ii) the brain states can be characterized as the oscillation with different spectral features and different spatial activations. The procedure for generating simulated MEG signals is depicted in Figure 2.

**Figure 2.**
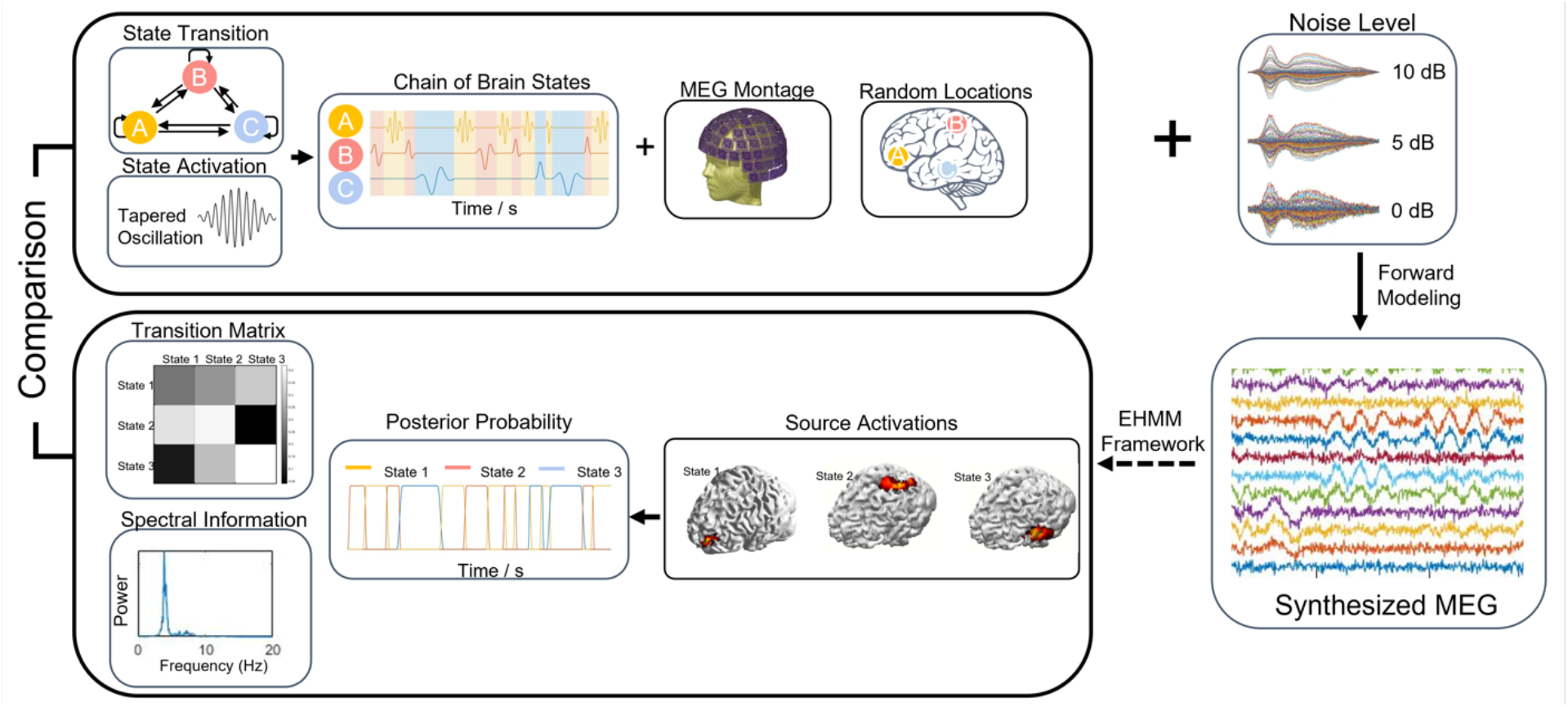
Simulation protocol overview. The chain of brain states was generated based on the state transition matrix and the state spatiotemporal profile. The chain of activation went through forward modeling to generate the synthesized MEG. Different levels of noise were added to the synthesized MEG. The MEG recordings were then put into the EHMM framework. The results were compared to the simulation parameters to verify the accuracy of EHMM model.

With the aforementioned assumption, we simulated each trial as a MEG segment with a few brain states, and each state was represented by an oscillation. For the spatial profile, Colin Brain template in Brainstorm toolbox was used as the head model (Tadel et al., 2011), and the cortical surface was segmented into 15,002 cortical vertices. For each brain state, we randomly selected one cortical point as the source location. In the cases of multiple sources, the distance between any two sources was set to be larger than 5 cm. For the temporal profile of each state, a frequency value was picked from 2-20 Hz to generate the temporal activations.

We first calculated the source level activations. The source level activation was simulated as multiple events where each event is one brain state activated for a certain length of time. To mimic a realistic case where the brain states are transitioning to each other, a Markov Model (Hidden Markov Model with Emission probability as an identity matrix) was used to simulate the time course. A transition matrix was then randomly generated as the probability of one state transitioning to the other one. The initial event was randomly drawn from all states and the following states were generated based on the transition matrix. Note that when one brain state is activated, i.e., when one region on the cortex is activated, the other regions remain silent. For each event, the activation time is randomly selected from 20-200 ms to represent the transient brain activities (Vidaurre et al., 2018). For a given length, the activation time course (oscillatory activity) was multiplied by a tapering Tukey window to mimic the state transition edge effect.

After obtaining the source level activation, we performed forward modeling to generate the synthesized E/MEG signal. While this technique is applicable on both EEG and MEG, we used a 102 channel Elekta MEG montage with magnetometers as the sensor configuration. A one-layer boundary element method head model was used to calculate the scalp MEG using the OpenMEEG toolbox (Gramfort et al., 2010) and the leadfield matrix was generated. Note that both three-layer BEM model (scalp, skull, and brain) and one-layer BEM model (brain only) are widely used in practice (Gramfort et al., 2010; Hallez et al., 2007; Tanaka and Stufflebeam, 2014). In this study, we adapted one-layer model since the neuromagnetic signals are minimally affected by the tissue conductivity (Hamalainen and Sarvas, 1989). A 2-min source activation was multiplied by the leadfield matrix to generate a 2-min MEG signal.

Different scenarios were simulated to evaluate the proposed framework. We varied the number of states from 4 to 7, with 4 being the most commonly used number for many state-space model as mentioned before. Also, Gaussian white noise was added to the generated signal to reach signal-to-noise ratio of 0 dB, 5 dB, and 10 dB to simulate the noise-contaminated MEG signals.

For each condition, 100 trials were simulated. The synthesized MEG recordings were fed into the EHMM framework. To be consistent with the literatures, a 100 ms embedding window was adapted and 50 initializations were used to obtain stable solution through empirical testing. Two types of result were then obtained from the EHMM framework: the spatiotemporal brain states profile represented by brief segments of MEG recordings, and the posterior probability for each of the states. The posterior probability was then put through the Viterbi algorithm (Eddy, 2004; Pulford, 2006) to get the most probable state activation.

### Simulation Evaluation

The result states were firstly compared to the simulated brain states to create 1-on-1 match by ranking the differences between all the pairs. To be more specific, all result brain states were compared to all simulated brain states, and the pair with smallest absolute difference was firstly selected, and the two matched states were removed from the pool. The next pair was then selected from the remaining values until each of the result states was matched to a simulated state. After doing so, for each trial, three evaluation metrics were obtained by comparing estimated states to its matching simulated brain state:

i. The result states were projected onto the source space using a Linearly Constrained Minimum Variance (LCMV) vector beamformer (Van Veen and Buckley, 1988; Woolrich et al., 2011). The Localization Errors (LEs) were calculated as the Euclidian distance from maximum of cortical source activation to the matching simulated brain state source location. LEs were averaged for all the brain states within each trial.
ii. A fast Fourier transform (FFT) was performed on each extracted state. The frequency with the highest power were extracted for each brain state to represent the temporal profile. The value was then compared to the simulated oscillation frequency for each state and the absolute difference was calculated. The absolute differences between all result states and simulated states were averaged.
iii. The correlation between result transition matrix and the simulated transition matrix in the 2-min MEG segments were calculated. For any timepoint *t*, if the current classified state *s*_t+1_ = *q_j_* was different from the state *s*_t+1_ = *q_j_*, a transition from state *g*_i_ to *q*_j_, was counted. We use *N*_i→j_ to denote the number of transitions from state *q*_i_ to *q*_j_ the transition probability matrix *Ā* is calculated as 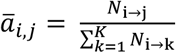.

In this case, we were neglecting the self-transition phenomenon where the brain state remains the same for two consecutive events. A smoothing procedure was introduced to detect the events with long durations and to break these events into smaller events. Note that the simulation transition matrix used to generate the MEG segment is slightly different from the actual simulated transition matrix, since the latter is a realization of the former based on Bayesian model, which could be different from trial to trial. In this simulation, we were comparing the result transition matrix to the latter, and the correlation coefficient (between 0 and 1) was obtained.

### Patient Data Acquisition and Preprocessing

The patients included in this study underwent MEG and intracranial EEG (iEEG) monitoring at the University of Pittsburgh Medical Center (UPMC) as a clinical routine. The data analysis study was approved by and performed in accordance with the regulations of the Institutional Review Boards (IRB) of Carnegie Mellon University and University of Pittsburgh.

A total of 10 focal drug-resistant epilepsy patients were included in this study. The patients were selected based on the following criteria: 1) Patients who underwent pre-surgical evaluation, including MEG recordings and MRI; 2) patients who underwent iEEG monitoring and the identified SOZ electrodes were located on the cortical regions (rather than subcortical locations such as amygdala or hippocampus); 3) patients who underwent resective surgery and had a postoperative follow-up of at least 12 months; 4) patients who had Engel I surgical outcome as rated by clinicians.

Each patient underwent a 306 channel recording using the Elekta MEG system (Elekta Neuromag, Helsinki, Finland) with 102 magnetometers and 204 planar gradiometers; a 10-minute run with magnetometers only (102 channels) was used in this study. The recorded MEG was band-pass filtered between 1-50 Hz. For each patient, an individual cortex surface model, a Desikan-Killiany (DK) atlas (Desikan et al., 2006), and a one-layer boundary element method (BEM) model were constructed from the pre-operational MRI. The sensors were co-registered using anatomical landmarks including Nasion, left pre-auricular point (LPA), right pre-auricular point (RPA), and digitized head points. The lead-field matrix was then calculated using the OpenMEEG (Gramfort et al., 2010) software in MATLAB. The preprocessing steps were all conducted with the Brainstorm software (Tadel et al., 2011).

### Brain State Extraction and Source Imaging

For each patient, the resting state data were visually inspected for removal of bad segments and bad channels. EOG and ECG channel was extracted from the original recording to remove EOG and ECG artifact. After that, the 10-min data were segmented into 2-min epochs and fed into the EHMM model to obtain the estimated brain states. For each epoch, two parameters, the 100 ms embedding window and 50 randomized initializations, were adapted from simulation.

After obtaining the states from the EHMM model, the brain states (sensor data) were projected onto the source space using a LCMV vector beamformer carried out separately on each state.

### Network Construction and Feature Index

Partial directed coherence (PDC) was used to estimate the directed functional interactions for each brain states. PDC is based on the concept of Granger-causality (Baccalá and Sameshima, 2001; He et al., 2019) and is computed using multivariate autoregressive (MVAR) models which simultaneously models multiple time series. To reduce the dimension for MVAR calculation and reduce spatial leakage, after obtaining the LCMV solution, we mapped each individual cortex to the 68-ROI DK brain atlas as described before and averaged the source activation for each region. Thus, we computed PDC using the source activity of the 68 regions. For each state, the connectivity matrix (ROIs × ROIs) represented the flow from one ROI to another, averaged over the frequency range (1-50Hz).

### Graph Feature Index for Brain States

With the PDC connectivity pattern constructed, we selected 20% largest values which corresponding to the major information flow in the network and obtained a weighted undirected network *G* = (*V, E*) consisting of a set of vertices *V* and a set of edges *E* between them. An edge *c* connects vertex *v*_i_ with vertex *v_j_*. In this study, two metrics were used to describe global network properties, namely the average clustering coefficient and the betweenness centrality difference.

Clustering coefficient (CC) is a characteristic parameter to describe local clustering features of a network. The clustering coefficient for a vertex is then given by a proportion of the number of links between the vertices within its neighborhood divided by the number of links that could possibly exist between them. For each node, a high CC value indicate tight connections to the neighboring nodes thus forming a local hub. In a small world network, i.e., the physiological brain network such as resting state networks where most of the nodes connected to their nearest neighbors, but a few of them can spread over a long range, average CC are relatively high. In the pathological network, or the epileptic network, one would assume that only a few nodes occupying the most outflow where most of the other nodes are “muted”, which would lead to a low averaged CC value.

Betweenness centrality (BC), on the other route, is a key metric that is used to identify important actors in a network. It is a popular graph analysis technique based on shortest path enumeration. This metric is widely used to identify the key nodes in the brain networks (Wilke et al., 2011). We averaged BC value in the left hemisphere and the right hemisphere, and the difference is used to represent the lateralization of the network. Higher difference between the two hemispheres indicates a more lateralized network while a lower difference value indicates a more balanced network. We also observed that betweenness centrality value differences follow a normal distribution, which further verified our assumption.

The two metrics are meant to evaluate the properties of individual brain states. From previous studies as well as experiences, a pathological state is usually corresponding to the abnormal strong cortical activations where one node is driving the other parts of the brain. With this assumption, the qualified pathological state should be corresponding to a smaller averaged clustering coefficient (ACC) value and a larger betweenness centrality difference (BCD) value, which corresponds to a more centralized hub as well as a lateralized driving spot. After empirical investigations, we designed the Graph Feature Index (GFI) as:

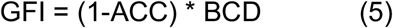

where a higher GFI represents higher probability to be a pathological state. The network analysis was performed using NetworkX toolbox implemented Python (Hagberg et al., 2008).

### Evaluation Metrics

Seizure onset electrodes identified from iEEG were extracted and marked by experienced epileptologists for each patient. Localization error was defined as the minimum distance between the source location with the maximum activation from the LCMV solution to the closest SOZ electrode, for each extracted state.

For each patient, five 2-min segments were analyzed, and all extracted brain states underwent a hierarchical clustering based on the connectivity patterns. The localization error was averaged within the cluster and the GFI is calculated for each of the grouped states based on the averaged connectivity pattern. The state with the highest GFI was identified as the pathological state for each patient.

### Interictal Epileptiform Discharges Analysis

Interictal Epileptiform Discharges were also analyzed to provide a baseline. For each patient, the MEG waveform was visually inspected to identify the IEDs. Two researchers must agree on the identified IEDs, for an identified event to be listed as an IED. While multiple types of IED were found in these patients, only IEDs showed concordant source localization results with the identified SOZ region were included for further analysis, as the purpose of this IED analysis was to provide a benchmark instead of investigating the diversity of IEDs. The IEDs identified were averaged and then fed into source imaging algorithm (LCMV) to obtain the source solution. The localization error was also calculated as the minimum distance between the source location with the maximum activation from the LCMV solution to the closest SOZ electrode.

## Results

### Simulation Results

As shown in Figure 3B, under various SNRs (0, 5, 10 dB) and various number of states (N = 4, 5, 6, 7), the EHMM provided robust estimates of source location overall. The localization error increases as the SNR decreases. This indicates that in the simulation setting, SNR could influence the EHMM’s performance largely, but the results remain reasonably stable even in extreme scenario where the signal-to-noise ratio is 0 dB. Additionally, the localization errors increase as the number of simulated brain states increases, but generally the localization error is less than or around 10 mm.

**Figure 3.**
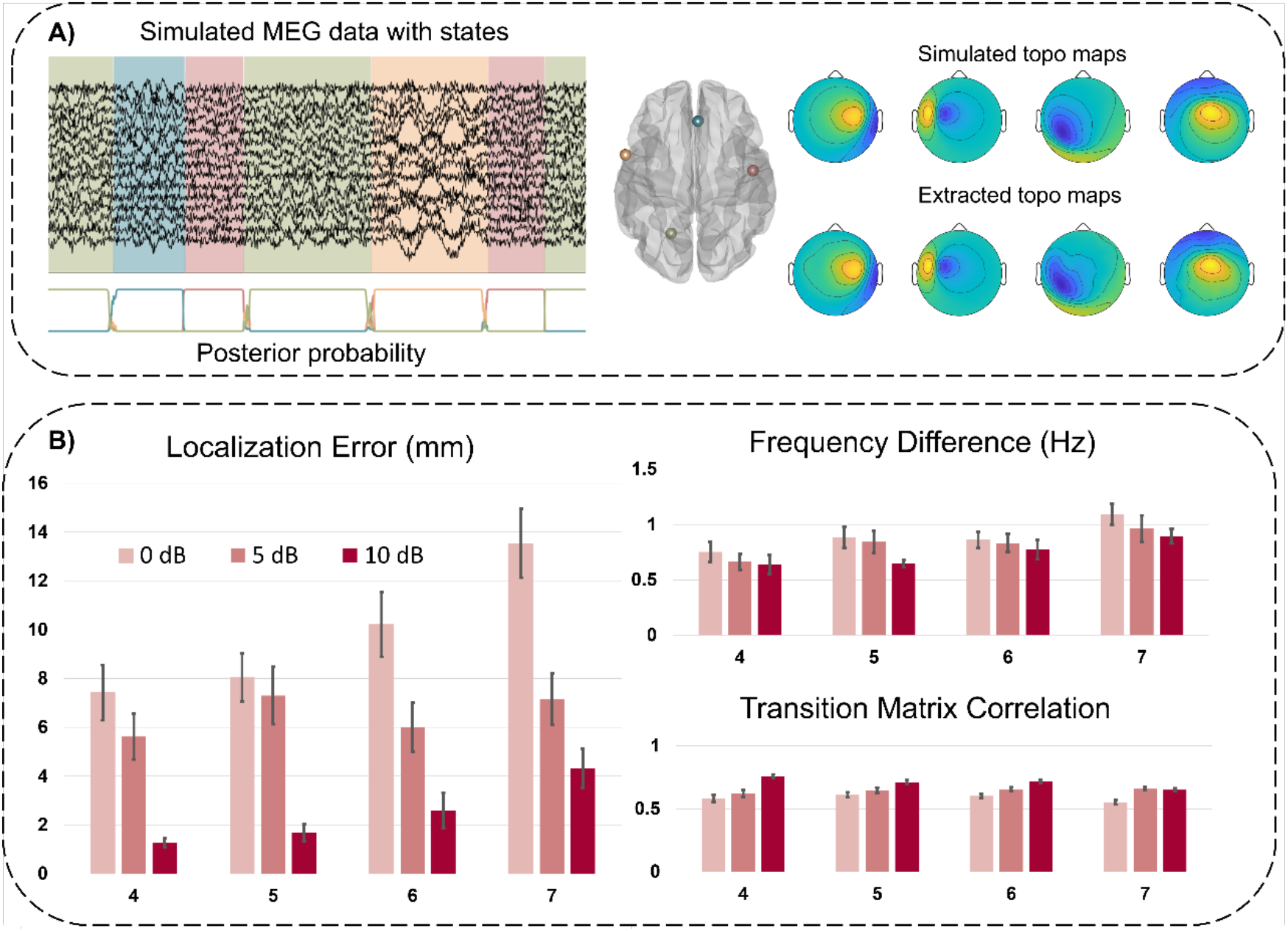
Simulation results. A) Example of one trial with 4 simulated states, represented by four different colors. The simulated MEG is overlaid with colors to represent the current simulated brain states. The EHMM framework successfully extracted the states, represented by the topo maps in sensor space. This trial is with 0 dB noise. B) Quantitative evaluations. Left panel: Localization error (in mm). The extracted states were projected to the cortex and the source region with maximum activation was compared to the simulated source (dipole model). For each trial, the average localization error was calculated for all the matched state pairs. The black bar indicates standard error. Right top panel: Frequency Difference (in Hz). The maximum frequency for each state was extracted and compared to the simulated frequency. Right bottom panel: transition matrix correlation. The correlation between simulated transition matrix and the result transition matrix was calculated for each trial.

From Figure 3B, it can also be seen that the frequency differences are small. The frequency differences represent the differences between the simulated oscillation and the obtained brain state temporal profile. The difference of approximately 1Hz indicates that the EHMM framework not only captured the spatial information but also the intrinsic oscillatory power of the brain states. Moreover, one of the features of HMM model over other clustering methods is the ability to accurately capture state transition, and this can be observed from the correlation between simulated transition matrix and the obtained transition matrix.

### Patient State Feature Extraction

One patient example is shown in Figure 4, where a right temporal SOZ was identified. The patient underwent left temporal lobectomy and was seizure-free at 1-year follow up. A segment of MEG time course is shown in Figure 4B where two extracted states were shown by the pale green boxes. The left-side state is showing a more balanced and spread-out connectivity, while the right-side state is showing a centralized activation from the right posterior temporal region. The representative topo map for the example patient can be found in Figure 4C, which shows a clear right lateral temporal dipole pattern.

**Figure 4.**
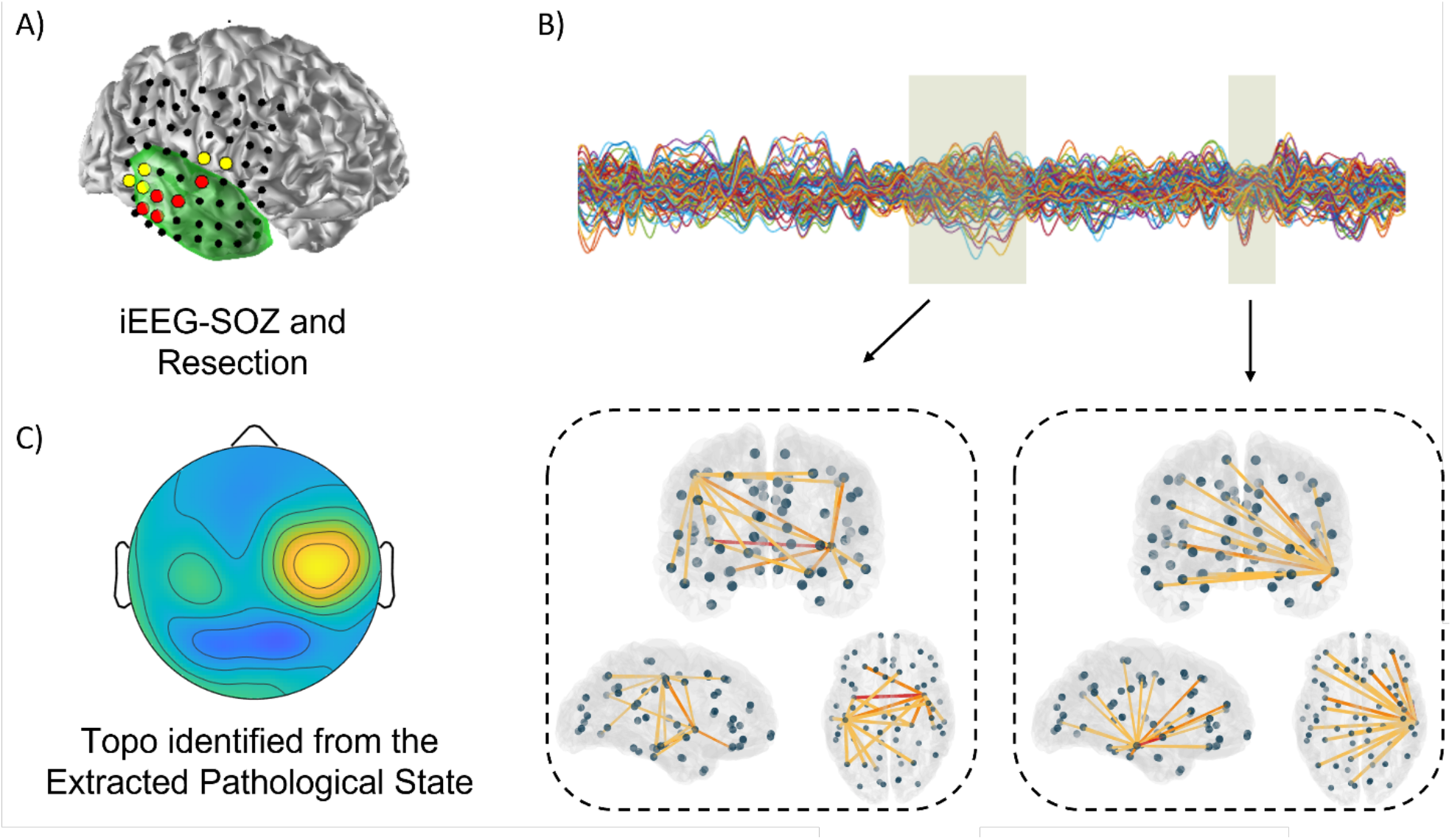
Patient Example. A) Example of one patient with right temporal epilepsy. Green color shows the resection and each dot represent one intracranial EEG electrode. The red color shows the seizure onset zone. The yellow color shows the seizure spread. B) A segment of MEG data and two example brain states. In each of the brain network figure, each grey dot represents the center of one ROI in the DK atlas. The red color edges represent the strongest connection between the ROIs (maximum PDC value). The orange color edges represent the strongest 5 connections. The yellow edges connecting dots represent the strongest 20 connections. The first one lasted longer and showed bilateral, spread-out network with local hubs. The latter one showed lateralized, centralized activation. The connectivity patterns were thresholded differently to better represent the network features. C) The topo map of the identified pathological state. The time point with highest global field potential is selected as the localization error is calculated in this time point.

In 10 patients studied, we extracted on average 6 different brain states (as shown in Figure 5A). We then calculated the Averaged Clustering Coefficient and the Betweenness Centrality Difference to get the Graph Feature Index for each state. The range of GFI varies from 0 to 0.03 as shown in Figure 5B. By contrasting the GFI value with the localization error for each state, the distribution of GFI can be classified into a spread-out points cluster in the left side, which is corresponding to a higher ACC with lower BCD states, or more likely the normal physiological brain networks. On the other side, points with GFI larger than 0.025 forms another cluster which coincidently related to an obvious smaller localization error.

**Figure 5.**
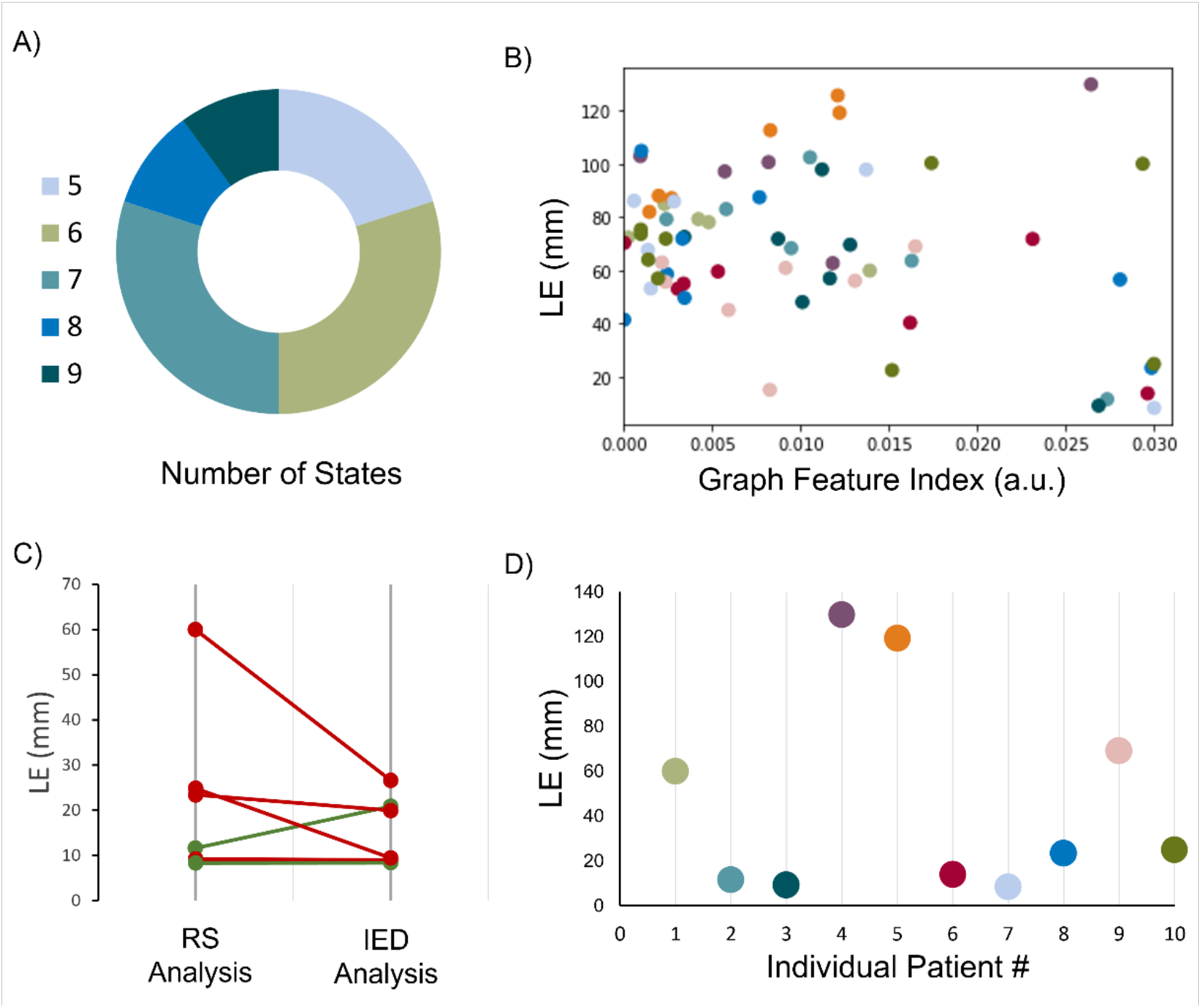
Pathological state analysis results. A) The number of extracted brain states for all patients. Majority of the patients had 6-7 brain states extracted; B) The Graph Feature Index for all the patients. Each color represents one patient. C) Resting state analysis comparing to the IED analysis, in 6 patients with concordant IED cluster. Red lines indicate worse results while green lines indicate better LE. D) Localization error for individual patient, by using the state with highest GFI to represent the pathological state. The colors are corresponding to the colors in B).

### Extracted Pathological States vs IED Analysis

If we select the state with largest GFI as the most pathological state, the individual patient outcome was then represented by this selected pathological state. For each state, we selected the time point with the highest energy to extract the source imaging results and compared to the iEEG-SOZ.

As shown in Figure 5D, 6/10 patients showed localization errors around 20 mm. These selected states are also well-captured in the right bottom corner of the Figure 5B. It is noteworthy that the maximum GFI for some patients are relatively small (the orange color, corresponding to patient #5).

In 6/10 patients, we extracted good amount of IEDs (> 5) and conducted the source localization on the averaged extracted IED. In these 6 patients, we compared the results from the extracted pathological state to the IED analysis results and showed comparable results in 5/6 patients as shown in Figure 5C. In one patient where no consensus can be reached using IED analysis, the resting state analysis still gave a 13.82 mm result (as shown in Figure 5D, patient #6, dark red point).

## Discussion

In this study, we have developed a framework to analyze the resting state E/MEG recordings for the purpose of identifying and stratifying the underlying brain processes. Our framework includes a two-step strategy. We firstly segmented the E/MEG signal into a few transient spatiotemporal brain states using an embedded Hidden Markov Model. After that, a graph-feature-index combining betweenness centrality difference and the average clustering coefficients was generated to evaluate the likelihood of a brain state to be considered as pathological state or not. The proposed framework was evaluated and verified in computer simulations. In 10 focal epilepsy patients who are seizure-free after resective surgery, we then applied the proposed framework on a 10-min MEG recording for each patient. The identified pathological state source localization results were compared to the SOZ identified from intracranial EEG recordings. In 6/10 patients, we obtained concordant results with the SOZ and obtained average localization error of 15.23 mm. In the subgroup of patients where interictal epileptiform discharges are abundant, the concordance rate is particularly high. It is noteworthy that while the validation was performed on epilepsy patients with MEG recordings, the methodology is applicable to both MEG and EEG.

Resting states analysis has been pursued from various angles. Several MEG or EEG analysis methods have been considered in epilepsy studies (Antonakakis et al., 2016; Rotondi et al., 2016), focusing on connectivity measures such as coherence analysis (Elisevich et al., 2011) and phase lag index analysis (Nissen et al., 2017, 2016) on coarsely-parcellated brain models. Topographical microstate analysis, as described before, is another emerging field, where a few typical patterns in raw EEG/MEG are extracted and compared to healthy subjects to identify patients from healthy controls (Khanna et al., 2015; Yuan et al., 2012). Although good accuracy can be obtained, especially combined with machine-learning methods (Ahmadi et al., 2020; V et al., 2018), the extracted states as well as their transitions are more representative in a group level instead of individual level. While some studies showed good accuracy in identifying epilepsy patients from healthy controls or lateralization in TLE patients (Coito et al., 2016; Nissen et al., 2017; Vollmar et al., 2018), the findings heavily rely on the patient groups and the control groups. Moreover, an epileptic seizure onset zone, as needed in the clinical diagnosis for further surgery planning, is still challenging to obtain through the aforementioned studies. The strength of our proposed framework, in this context, is that we can extract the brain states and identify the pathological state for individual patient without the requirement of a control group as the baseline. More specifically, the majority of the physiological states in MEG recording would be the normal “baseline” activity of the brain while the abnormal activations would stand out and be detected through our proposed framework, thus potentially indicating where the epileptic source is.

Due to the complex nature of the brain networks especially in epileptic brain, the physiological or the normal brain states and the pathological or the epileptic brain states could be mixed. Our solution to this problem was two-fold. We first followed the state-space model and adapted the EHMM where we make the most use of the high temporal resolution of E/MEG recordings to capture transient spatio-temporal dynamics and verified with simulation. In real patient analysis, even with disentangled brain states, the feature of a so-called “pathological state” is not yet fully understood or fully investigated. The underlying pathology of seizure generation most likely involves both abnormal brain structures and aberrant connections among these regions, leading to skewed large-scale network phenomena (Engel et al., 2013). In this work, we did not exclude the interictal epileptiform discharges from the original recording to create the so-called “spike-free” resting state recording (Grouiller et al., 2011). In practice, interrater reliability is still a challenging issue for correct identification of IEDs (Bagheri et al., 2017; Jing et al., 2020), and multiple types of IEDs co-existing in the interictal data also contribute to the complexity (Ye et al., 2021). Excluding IEDs, on one hand, may demonstrate the power of such analysis, while on the other hand, could be biased towards the IEDs selected to remove. Moreover, since the essence of HMM is dependent on the state transition (Eddy, 2004), removal of an important component from the recording could potentially jeopardize the integrity of the algorithm itself. By including entire segments of data, we treat the IED-related network as a part of the brain states and relies on the EHMM to separate the states from others. The abnormality of the IED networks as studied previously (Costa et al., 2021; Erem et al., 2015) could then contribute to identify such abnormality from the normal resting state network.

We then focused on application of graph theory, which provides a mathematical framework to characterize different topological properties of a network’s organization, and has been applied on various data format to aid the presurgical evaluation (Case et al., 2019; Wilke et al., 2011). In this work, we looked at two fundamental concepts in graph theory, segregation (i.e., local connectivity) and integration (i.e., global network functioning). Centrality is a key measure of integration. We calculated the averaged clustering coefficient to reflect overall properties of the network. A higher ACC is related to a more connected network with tight local connection, which is often seen in many of the large-scale physiological whole-brain networks. Betweenness centrality is a measurement of segregation. BC is computed based on the fraction of all shortest paths in the network that contain a given node. In other words, BC reflects the number of shortest paths from all nodes to all others that have to pass through a specific node. As such, a higher value of BC reflects the hubness of a node as an important, highly integrated influencer in the network. We used BC to determine the lateralization of the source, where differences of BC value between the left and right hemisphere were calculated to represent how lateralized the network connection is. It was observed in our data that majority of the extracted brain states show a high ACC value, and the BCD value follows a normal distribution, which aligns with prior whole-brain network studies conducted on healthy patients or epilepsy patients (Baker et al., 2014; Farahani et al., 2019; García-Prieto et al., 2017). An epileptiform-related state, by comparing the topography to the IED, showed a low ACC value and high BCD value (García-Prieto et al., 2017; Smit et al., 2008). One possible explanation is that a pathological state tends to have the focused energy around the seizure onset zone to facilitate the emission of IED or seizures (Bagheri et al., 2017; Costa et al., 2021). The connectivity is likely to increase in the SOZ and potentially in regions far from the SOZ, but to decrease for SOZ neighbors, leading to a global reduction of strength, leading to a decreased ACC. Thus, centralized regions tend to strengthen their connections with other hub regions but not so with the rest of the brain, leading to an increased BCD. One thing to note here is that while in some cases, brain networks extracted can be correlated with the healthy brain networks, we do not tend to build the one-on-one bonds, since the epileptic brain could be very different from the normal ones even if we performed a solid segmentation.

With the proposed resting state analysis framework, we were able to achieve a 60% accurate extraction of pathological states with a 15 mm localization error, while in some patients the GFI is not giving us the best results. It was also observed that in those patients with inaccurate results, even though we used the state with the highest GFI to represent the putative pathological results, the highest GFI is still much smaller than that in the other patients. Combined with the fact that we were not able to find a concordant IED cluster with a good amount of IEDs in 4/10 patients, this might indicate that the data is not long enough for the proposed framework to separate a valid abnormal pattern, or the pathological states were potentially blurred by the noise due to the non-invasive recordings. While such a caveat is inevitable in data-driven framework, it should be noted that such cases would be challenging for clinicians as well. Moreover, in one patient where standard IED extraction failed to lead to any conclusive results, we extracted a pathological state with high accuracy. In a word, the analysis was performed on a 10-min MEG recording and the pathological states were extracted in a data-driven manner, which could provide an overview of the data for clinical diagnosis and as a potential pre-filtering process to identify states/intervals of interest for the clinical team. The value of the proposed framework is particularly high when longer recordings are available, where manual search is not possible or practical.

Nonetheless, this approach also suffers from various technical limitations. While the PDC connectivity and the network features are both widely used in many other literatures, much more potential parameters could be tested to improve the framework (Malinowska et al., 2014; Mantini et al., 2011; Molen, 2016; Niso et al., 2015). Several studies have adapted more advanced machine learning algorithms to test for the best combination of parameters (Jin and Chung, 2017; V et al., 2018; Wu et al., 2020), which should be considered for future studies. We included only patients who had a good surgical outcome or in whom the focus was confidently localized with source imaging on IEDs. Future studies could explore the prediction of different degrees of non–seizure freedom. Along these lines, future studies should also contemplate the changes that occur in different segments of the data, which has been clustered by hierarchical clustering method in this framework due to the short duration of the MEG recording (10 minutes). In longer recordings where the distribution and the involvement of brain networks could vary and the occurrence of the pathological state could vary (Karoly et al., 2016), a more curated framework with sliding windows could be more informative in evaluating the different stages of the resting state data. Transition matrix, as an important feature of the Hidden Markov Model, was not fully investigated due to the short duration of the data and the relatively small patient population. Future studies could look into mechanisms of the pathological states and interactions with the other brain networks for a further understanding of the epileptogenesis. Despite all these impediments, our results promise an alternative way to identifying and stratifying epileptic networks from resting-state recordings.

To summarize, in this study, we have proposed a framework to extract pathological state from resting state electromagnetic recordings and achieved reasonable accuracy. Our results indicate that the brain networks can be disentangled from the resting-state electromagnetic recording and could be identified based on the connectivity features. The data-drive framework requires minimal human intervention and could potentially guide the surgical intervention for focal epilepsy patients undergoing presurgical planning. Given the generalizability of state space brain network model and increasing studies working on network features of other neurological disorders or even healthy brains, the proposed framework may have applications to study other brain diseases or healthy brains.

## Acknowledgment

We are grateful to Abbas Sohrabpour, Zhengxiang Cai, Rui Sun, Xiyuan Jiang, Haiteng Jiang, and Jason Yang for useful discussions on data analysis. We thank participating patients and their families whose involvement and sacrifice made this work possible. We also acknowledge selfless dedication and invaluable efforts of the University of Pittsburgh Comprehensive Epilepsy Center (UPCEC) team and particularly the staff of the UPMC Presbyterian University Hospital (PUH) Epilepsy Monitoring Unit (EMU).

